# Contextual associations of real-world objects guide attention in visual search

**DOI:** 10.64898/2025.12.12.693961

**Authors:** Lu-Chun Yeh, Belma Seferovic, Marius V. Peelen, Daniel Kaiser

## Abstract

Growing evidence indicates that visual search is influenced not only by perceptual but also by semantic information. Among such semantic influences, categorical effects have been extensively studied. By contrast, the influence of contextual associations, that is, how objects commonly co-occur within specific scene contexts, remains less understood. Here, we orthogonally manipulated contextual and categorical relationships between target and distractor objects in a visual search task, while controlling for perceptual target-distractor similarity. By applying linear regression analyses, we modelled behavioral responses and event-related potentials (ERPs) from contextual, categorical, and perceptual target-distractor similarity. We found that contextual associations guide attention by interacting with categorical information (modulating the N2pc) and ultimately shape behavioral search performance jointly with perceptual and categorical similarity. These findings demonstrate that contextual object associations can guide attention, thereby supporting efficient search in complex visual environments.

## Introduction

Visual search is a critical task during everyday life. When you have breakfast, for instance, you might look for a spoon to eat your morning cereal. Finding a spoon among other pieces of cutlery in a drawer often takes longer than locating it on the kitchen counter, since all cutlery shares similar perceptual features (e.g., silver color, elongated shape). Many studies indeed show that search performance systematically declines as perceptual similarity between targets and distractors increases (Duncan & Humphreys, 1989; Proulx & Egeth, 2006; Alexander & Zelinsky, 2012; Barras & Kerzel, 2017; Yeh et al., 2019). Theories of visual search suggest that such effects are mediated by search templates that bias visual processing and guide attention toward likely target features (Duncan & Humphreys, 1989; Desimone & Duncan, 1995; Wolfe, 1994, 2007, 2021). When distractors share features with the template, they divert attention away from the target, leading to poorer search performance.

However, real-world objects are not just defined by perceptual features but also by semantic features. Increasing evidence demonstrates that semantic properties also play a crucial role in guiding attention and shaping search performance. When targets and distractors share category membership (e.g., animacy), search performance is impaired (Belke et al., 2008; Telling et al., 2010; Wyble et al., 2013; Seidl-Rathkopf et al., 2015; Yeh & Peelen, 2022; Bahle et al., 2025). Complementing these behavioral effects, EEG studies showed that target-distracter similarity in semantic features (particularly object category) diminishes the N2pc component, a neural correlate of efficient attentional allocation (Telling et al., 2010; Yeh & Peelen, 2022), and that semantically related distractors can elicit an N2pc when the target is absent (Nako et al., 2014; Nako, Wu, & Eimer, 2014). These findings suggest that object categories, like perceptual features, guide the allocation of attention during visual search. Yet, categorical and perceptual similarities interact hierarchically during search: Attention is initially guided by perceptual properties and subsequently by categorical information when perceptual cues are insufficient (Yeh & Peelen, 2022).

Given the focus on category effects, other semantic influences have not been studied sufficiently. Object semantics encompasses not only categorical relationships but also contextual associations (Mirman et al., 2017; Nah & Geng, 2022). Contextual associations reflect the typical co-occurrence of objects within a specific scene context (e.g., cutlery and food commonly appear together in a kitchen). Such contextual associations facilitate object recognition when objects appear in a predictable scene context (see reviews: Bar, 2004; Oliva & Torralba, 2007). Yet, contextual associations are also activated automatically during object perception (Biederman, 1972; Cornelissen & Võ, 2017; Nah et al., 2021), supported by “context frames” that store these associations in memory (Bar, 2004). Recent neuroimaging studies revealed that even objects in isolation activate representations of contextually related objects in the visual system (Bonner & Epstein, 2021; Yeh et al., 2026). This automatic co-activation of contextually related objects prompts the hypothesis that contextual associations effectively guide visual search. If this hypothesis is true, it further begs the question of whether contextual similarity guides search in similar or different ways compared to perceptual and categorical similarity.

To address these questions, we investigated how contextual, categorical, and perceptual similarity independently and in concert influence visual search. We orthogonally manipulated the contextual and categorical relationships between targets and distractors. We additionally quantified continuous perceptual target-distractor similarity for each stimulus pair using an independent same–different task (Jacob & Arun, 2020; Yeh & Peelen, 2022; Yeh et al., 2026). We then applied linear regression analyses to behavioral responses and EEG data to quantify the effects of contextual, categorical, and perceptual similarity, as well as their interactions. To examine how contextual associations influence attentional guidance during search, our EEG analyses focused on the N2pc, an ERP lateralized component that usually emerges approximately 200–300 ms after stimulus onset, respectively, and over posterior scalp sites. The N2pc is characterized by greater negative amplitudes contralateral to the target field compared to the ipsilateral side; a stronger contralateral–ipsilateral difference reflects greater attentional allocation to the target (Luck & Hillyard, 1994; Eimer, 1996). If contextual associations indeed guide attention during visual search, distractors from the same scene context should draw attention away from the target, leading to a reduction in N2pc amplitudes as well as poorer search performance.

## General Method

### Participants

We conducted two experiments. Experiment 1 was a pilot study, and Experiment 2 was the main study.

In Experiment 1, sixteen healthy volunteers (14 females, 4 males, age = 23.56 ± 3.52 years) participated. This sample size followed a similar previous study (Yeh & Peelen, 2019).

In Experiment 2, twenty-five healthy volunteers (18 females, 7 males, age = 25.00 ± 3.12 years) participated. One participant did not complete the experiment due to a technical problem during EEG-acquisition, resulting in 24 datasets included in the final analysis. For this experiment, the required sample size was estimated from a power analysis based on the results of Experiment 1. The analysis showed that at least 20 participants were needed to yield 80 % power to detect the behavioral contextual × categorical similarity interaction observed in Experiment 1 (Cohen’s d = 0.59).

In both experiments, participants were native German speakers and had normal or corrected-to-normal vision. They provided written informed consent prior to participation and were compensated at a rate of 10 euros per hour. The study was approved by the Ethics Committee of Justus Liebig University Giessen and conducted in accordance with the 6th revision of the Declaration of Helsinki.

### Stimuli

We used a set of 24 real-world objects, which were also used in our previous neuroimaging study (Yeh et al., 2026). The stimuli stemmed from two contexts (12 kitchen objects and 12 garden objects) and two categories (12 tools and 12 non-tools). These objects were grouped into six sets of four items each, each of which contained one object per condition (i.e., 1 kitchen tool, 1 garden tool, 1 kitchen non-tool, and 1 garden non-tool). Within each set, objects were matched for their overall shape (see Fig. 1A). There were four exemplars per object, yielding 96 unique stimuli in total. All images were converted to grayscale, fitted into an invisible frame of approximately 4° × 4° visual angle, and presented on a white background. For an image-based analysis of luminance and perceptual similarity, see Yeh et al. (2026).

**Figure 1.**
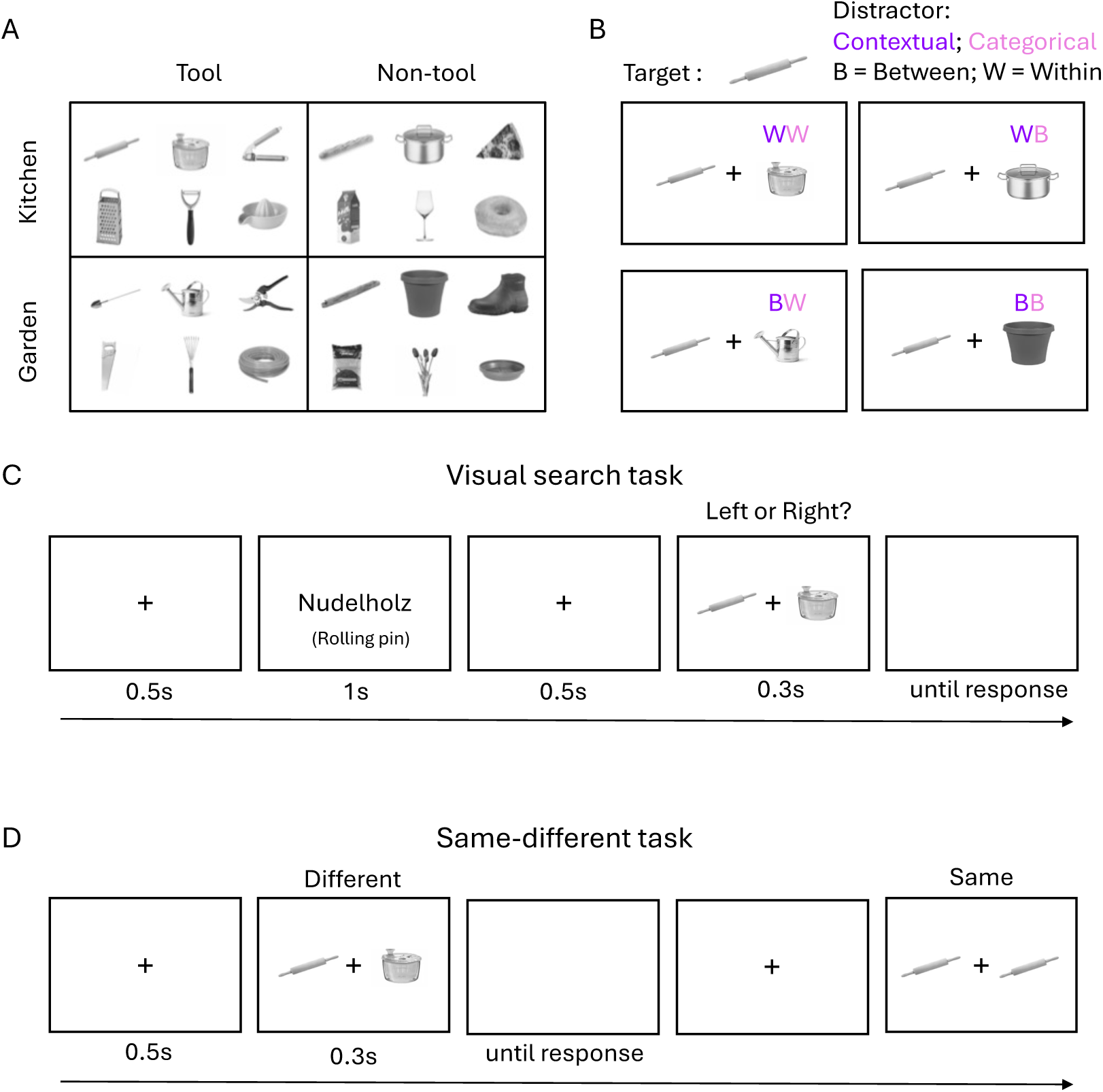
Stimulus, design, and task procedure. (A) Stimulus set. 24 objects were used in the experiments, selected from two contexts (kitchen and garden) and two categories (tool and non-tool). (B) A 2 (contextual similarity: between, within) X 2 (categorical similarity: between, within) within-subjects factorial design was employed. For each target object (e.g., a rolling pin), four types of distractors were presented together in a search array. (C) Visual search task. Participants were instructed to search for a predefined target (in German) within a search array and press the button corresponding to the target location. The English cue shown here was not presented in the actual experiment. The maximum blank response window was 0.8 s, followed by a blank intertrial interval that varied randomly between 0.8, 1.0, and 1.2 s. (D) Same-different task. Participants indicated whether two simultaneously presented objects were the same or different.

### Experimental Design and Paradigm

The visual search experiment employed a 2 (contextual similarity: between, within) X 2 (categorical similarity: between, within) within-subjects factorial design. Each object was paired with the other 23 objects, yielding 276 stimulus pairs. In each pair, one object served as the target and the other as the distractor. For contextual similarity, pairs were drawn from the same context (both kitchen or both garden; within-context) or from different contexts (one kitchen, one garden; between-context). For categorical similarity, pairs were either both tools or both non-tools (within-category) or a tool and a non-tool (between-category). The example pairs for each condition are illustrated in Figure 1B. Each condition contained 72 pairs, except the within-context/within-category condition, which included 60 pairs.

At the beginning of each trial, a fixation cross appeared at the center of the screen for 500 ms to signal trial onset. Next, a target word cue (presented in German) was shown for 1000 ms, followed by another 500-ms fixation. A search display containing two objects was then presented for 300 ms and subsequently replaced by a blank screen for up to 800 ms, during which participants could respond. Their task was to indicate the location of the target. Responses were made bimanually, using the index fingers to press either the F key for a left-side target or the J key for a right-side target. Target locations were fully counterbalanced across trials. The intertrial interval varied randomly between 800, 1000, and 1200 ms. The overall procedure is illustrated in Figure 1C.

### General Experimental Procedure

In both experiments, participants first completed a questionnaire to assess their knowledge about the 24 selected objects. Here, the (German) words representing all objects were presented, and participants were asked whether they knew the object. If a participant did not know an object, the experimenter described it to ensure accurate identification during the task. After that, participants practiced a block of 12 trials showing the four different objects not used in the formal experiment (spoon, mug, sickle, birdhouse). During the formal experiment, participants were required to fixate on a cross at the center of the screen and respond as quickly and accurately as they could. The experiment consisted of 12 blocks, each containing 92 trials, resulting in a total of 1,104 trials. There were four trials per stimulus pair, and each stimulus appeared as a target twice. Accordingly, each condition comprised 288 trials, except for the within-context/within-category condition, which included 240 trials. Within each block, all conditions were represented: 24 trials per condition, except for the within-context/within-category condition, which had 20 trials (the same object was never shown in both locations). Target identity (which stimulus served as the target) and target position (left vs. right) were counterbalanced across blocks. Stimulus pairings, including the use of different exemplars, were also counterbalanced across blocks. Trial order within each block was randomized. Participants could rest between blocks and initiate the next block at their own pace. The total duration of the experiment was approximately 60 minutes per participant. In Experiment 2, EEG signals were recorded while the experiment was performed, and the participants were additionally asked to blink only after making a response.

After completing the search task, participants filled out a co-occurrence questionnaire covering all object pairs, designed to validate the assignment of objects to the two contexts. The questionnaire comprised two parts: (1) object–scene co-occurrence and (2) object–object co-occurrence. In the first part, participants indicated the typical context for each object (kitchen or garden). In the second part, they rated, on a scale from 0 to 100, how often each object pair co-occurs in daily life and how similar their functions are—that is, whether the objects are used in comparable ways for similar purposes. Across both experiments, the selected objects were consistently rated as being more frequently encountered in their assigned context, and object co-occurrence was reliably lower in the between-context condition than in the within-context condition (Between = 5.84, Within = 38.67, *t*[39] = 15.27, *p* < .001).

### EEG recording and preprocessing

Electrophysiological data were recorded using an Easycap system with 32 channels and a Brain Products amplifier with a 1,000 Hz sampling rate. The electrodes included four sites in the central line (Fz, Cz, Pz, and Oz) and 14 sites over the left and right hemispheres (FP1/FP2, F3/F4, F7/F8, FC1/FC2, FT5/FT6, FT9/FT10, C3/C4, T7/T8, CP1/CP2, CP5/CP6, TP9/TP10, P3/P4, P7/P8, and O1/O2). AFz served as the ground electrode, and Fz served as a reference electrode. FP1/2 and FT9/10 were remounted to vertical and horizontal eye movement recorders and removed from the subsequent EEG analysis. The impedance of all the electrodes was kept below 20 kΩ. Triggers were sent from the presentation computer to the EEG computer via a parallel port.

Offline EEG data were preprocessed using the Fieldtrip toolbox (Oostenveld et al., 2011) in MATLAB (MathWorks). No filter was applied during recording and preprocessing. EEG data were re-referenced using the average of all electrodes and then epoched from −100 to 500 ms relative to stimulus onset. Epochs were baseline-corrected using the prestimulus period (from 100 to 0 ms). Epochs containing artifacts (i.e., blinks or eye movements) were rejected based on visual inspection. Epochs with incorrect responses were also excluded in further analyses.

### Perceptual similarity measurement task

To disentangle contextual and categorical effects from perceptual effects, an independent same-different task (N = 34) was conducted (see Yeh et al., 2026 for details). In this task, participants judged whether two simultaneously presented object images were identical or different (Figure 1D). RTs from the different-object trials, averaged across participants, were used as a continuous index of perceptual similarity, with longer RTs reflecting higher similarity (Jacob & Arun, 2020; Yeh & Peelen, 2022).

### Behavioral analyses

Behavioral data were analyzed using a linear regression model. Both accuracy and response times (RTs) were examined, with RT analyses restricted to correct trials. We first calculated the percentage of error and RTs from each stimulus pair across all exemplars and repeated trials, and constructed two 24-by-24 matrices. Next, to model behavioral performance, we constructed three predictor representational similarity matrices (RSMs): (1) a contextual RSM, coding whether object pairs originated from the same context (e.g., both kitchen or both garden) or different contexts (one from kitchen and one from garden); (2) a categorical RSM, coding whether objects belonged to the same category (e.g., both tools or both non-tools) or different categories (one tool and one non-tool); and (3) a perceptual RSM, derived from a same–different task (see above). The perceptual RSM was based on the group-averaged RT data (with perceptually more similar objects resulting in longer RTs). All RSMs were vectorized by retaining only the lower off-diagonal values. Behavioral performance vectors were then predicted using a linear regression model that included the contextual, categorical, and perceptual RSMs along with their interaction terms (Figure 2A). Beta estimates from the model were tested against zero using one-tailed one-sample t-tests, and false discovery rate (FDR) correction was applied to control for multiple comparisons.

**Figure 2.**
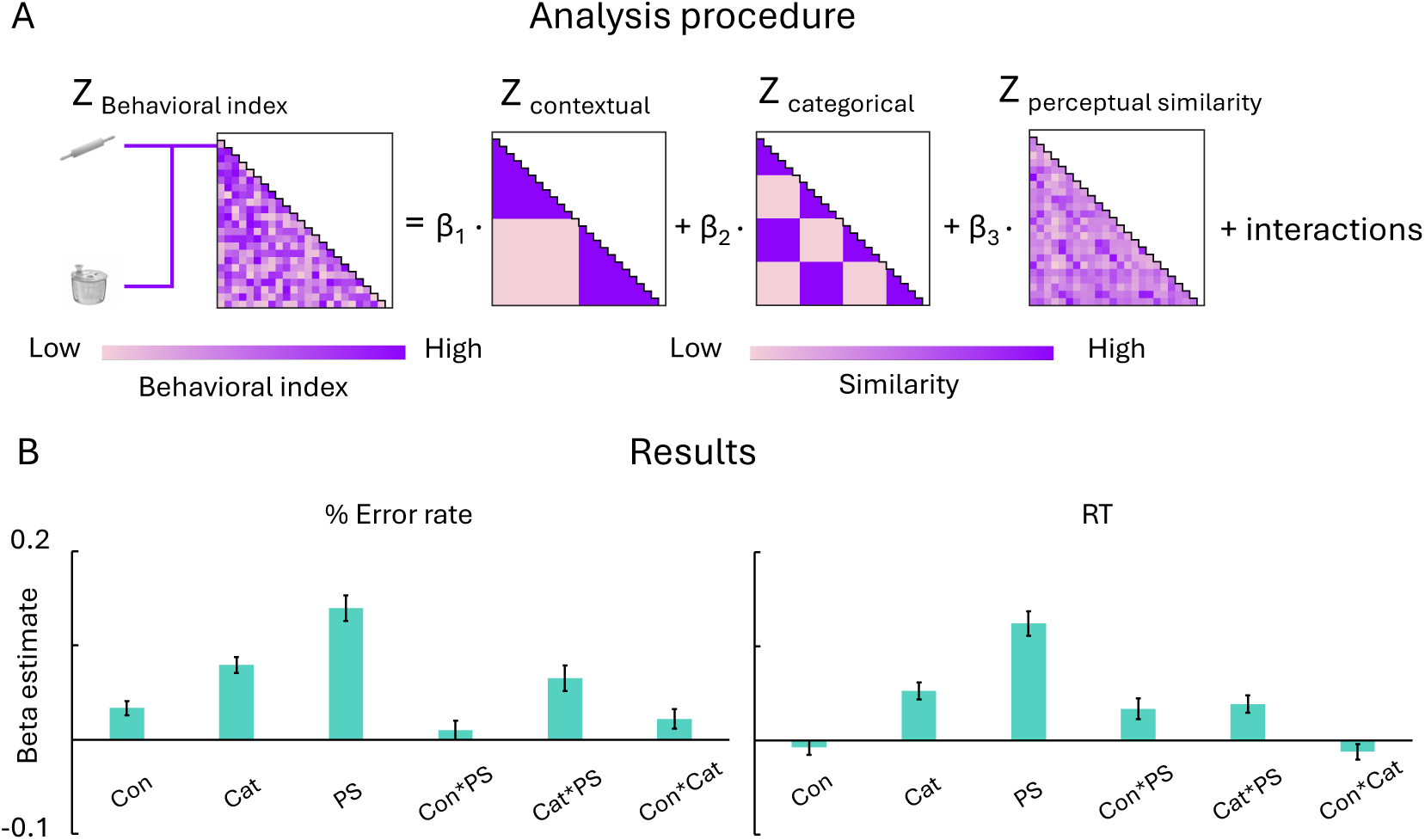
Behavioral analysis approach and results. (A) Analysis approach. Behavioral performance index matrices (computed from error rates or response times) were predicted from three model RSMs reflecting the objects’ similarities in (1) contextual similarity, (2) categorical similarity, (3) perceptual similarity, and (4-6) their pairwise two-way interaction terms. For each predictor, we estimated its beta weight in a linear model, separately for each participant. These beta weights indicate the contribution of contextual, categorical, and perceptual factors to search performance. (B) Pooled behavioral results for error rates (left panel) and response times (right panel) across both experiments. Error bars represent the standard error of the mean.

### ERP analyses

The effect of the N2pc component was quantified as the mean voltage within a specified latency window among three pairs of posterior electrodes, including P3/4, P7/8, and O1/2 (note that PO7/8 were not present in our EEG system). First, we examined the peak latency of the N2 waves across conditions and participants and defined 50-ms time windows centered on the peak latency (Luck & Hillyard, 1994). Next, contralateral-minus-ipsilateral difference waveforms were calculated relative to the target location for each trial. The resulting amplitudes were averaged within the predefined time window, where increasingly negative values represent a greater N2pc magnitude. Then, trial-based linear regression model analyses were applied. We fit a linear regression model to the z-transformed N2pc using z-transformed contextual, categorical, and perceptual similarity, and their interactions at the individual-participant level (Figure 3B). Finally, we tested beta estimates against zero using one-sample t-tests at the group level with FDR corrections.

**Figure 3.**
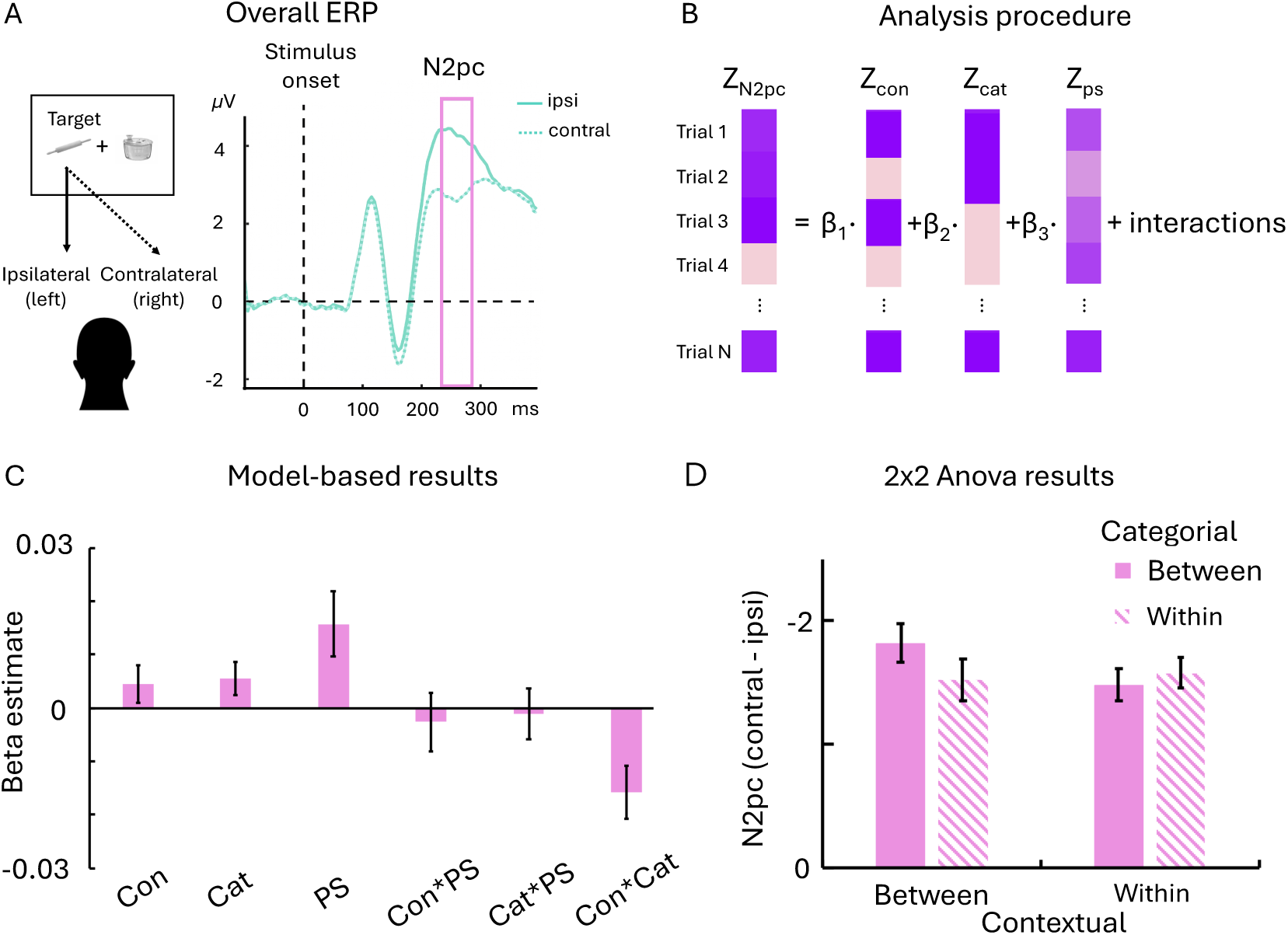
ERP analysis approach and results. (A) ERP waveforms averaged across all conditions and all participants over posterior electrodes (averaging electrodes P3/4, P7/8, and O1/2; ipsilateral waveforms: solid lines; contralateral waveforms: dashed lines). The purple/pink frame indicates the selected time window for the N2pc effect, a 50-ms window centered on the N2 peak latency. (B) Illustration of trial-based regression analysis. β represent the standardized slope of the predictors, including contextual, categorical, and perceptual similarity and their pairwise two-way interaction terms, resulting in six betas. (C) Results from the linear models for the N2pc, Con: contextual, Cat: categorical, PS: perceptual similarity. Error bars represent the standard error of the mean. (D) Results of a 2 (contextual) × 2 (categorical) within-subject ANOVA for the N2pc. Error bars represent the standard error of the mean.

## Result

### Behavioral results

We conducted linear regression analyses to disentangle contextual, categorical, and perceptual similarity effects and their interactions. As no three-way interaction was observed, the final models included only the three main effects and their pairwise interaction terms. All p-values reported here were adjusted by FDR corrections among 6 tests (three main effects and three two-way interactions). The analysis approach and behavioral results are shown in Figure 2.

First, we examined whether results differed across the two experiments using a mixed-design analysis of variance (ANOVA) with one between- (Experiment 1 vs. 2) and one within-subjects factor (six beta estimates). Type III sum of squares and Greenhouse-Geisser adjustment were applied to correct the unbalanced design of the between-subjects factor and the violation of sphericity. No between-subject effect or interaction was found for either error rates or RTs, indicating that the results from the two experiments revealed a consistent pattern (error: group effect, F [1,38] = 3.04, *p* = .089; interaction: F [3.825,145.349] = 0.09, *p* = .084; RTs: group effect, F [1,38] = 0.48, *p* = .493; interaction: F [5,190] = 1.02, *p* = .408). The two experiments were therefore pooled (N = 40) and analyzed together (see Supplementary Materials for separate analyses for Experiments 1 and 2).

The error rate results showed significantly positive beta estimates for contextual similarity (b = .034, t[39] = 4.33, FDR-corrected *p* <.001), categorical similarity (b = .080, t[39] = 9.24, FDR-corrected *p* <.001), and perceptual similarity (b = 0.140, t[39] = 10.21, FDR-corrected *p* <.001), indicating that the higher the similarity, the worse the search accuracy. We also observed two significant interactions. The interaction between contextual and categorical similarity (b = .022, t[39] = 2.09, FDR-corrected *p* =.026) indicates that the contextual similarity effect was stronger when categorical similarity increased, and the interaction between categorical and perceptual similarity (b = .065, t[39] = 4.83, FDR-corrected *p* <.001) indicates that the categorical similarity effect was stronger when perceptual similarity increased. No interaction between contextual and perceptual similarity was found (b = .010, t[39] = 0.98, FDR-corrected *p* =.165). The response time results showed significantly positive beta estimates for categorical similarity (b = .052, t[39] = 5.71, FDR-corrected *p* <.001) and perceptual similarity (b = 0.124, t[39] = 9.52, FDR-corrected *p* <.001), indicating that the higher the similarity, the slower the response. No significant contextual effect was found (b = −.007, t[39] = −0.98, FDR-corrected *p* =.922). We also observed significant interactions between contextual and perceptual similarity (b = .034, t[39] = 3.01, FDR-corrected *p* =.003) and between categorical and perceptual similarity (b = .038, t[39] = 4.09, FDR-corrected *p* <.001), indicating that the contextual and categorical similarity effects were stronger when perceptual similarity increased. No significant interaction between contextual and categorical similarity was found (b = −.012, t[39] = −1.45, FDR-corrected *p* =.922).

Together, the behavioral results reveal that contextual associations influence search performance both independently and in conjunction with perceptual and categorical similarity. However, behavioral data alone are insufficient to determine whether and how contextual associations guide attention during search, as they cannot disentangle effects on attentional allocation from those on later decision-making stages. Therefore, we next examined how contextual associations modulate neural markers of attentional allocation, specifically the N2pc ERP components.

### N2pc results

We first defined the N2 time windows based on the group data across three pairs of posterior electrodes. Across conditions and participants, the N2 peak was observed at 260 ms after stimulus onset. We analyzed N2 amplitudes by assessing the average voltage within 50ms windows centered on these peak latencies. We observed significantly more negative amplitudes in electrodes contralateral to the target, compared to ipsilateral electrodes, co-ip = −0.87, t[23] = −7.74, *p* <.001 (Figure 3A). We thus quantified the N2pc within the 235–285 ms window, for each trial and participant. Trial-based linear regression analyses (Figure 3B) were then performed to assess the three similarity effects and their interactions. Consistent with the behavioral results, no three-way interaction was observed. Therefore, the final models included only the three main effects and their pairwise interaction terms. All p-values reported here were FDR-corrected across 6 tests (three main effects and three two-way interactions).

##Figure 3C shows the results of the linear regression analyses on the N2pc data. N2pc results only showed a significantly positive beta estimate for perceptual similarity (b = .016, t[23] = 2.58, FDR-corrected *p* =.025). This positive relationship indicates that greater perceptual similarity between the target and distractor predicted a more positive contralateral-minus-ipsilateral magnitude (i.e., smaller N2pc). In contrast, no significant effects were found for contextual similarity (b = .0045, t[23] = 1.27, FDR-corrected *p* =.161) or categorical similarity (b = .0055, t[23] = 1.74, FDR-corrected *p* =.096). No significant interactions between perceptual similarity and either contextual similarity (b = −.0026, t[23] = −0.48, FDR-corrected *p* =.683) or categorical similarity (b = −.0011, t[23] = −0.22, FDR-corrected *p* =.683) were found. However, we observed a significantly negative beta estimate for the interaction between contextual and categorical similarity (b = −.0158, t[23] = −3.14, FDR-corrected *p* =.013, left-tailed).

To further examine this interaction, we then analyzed our ERP data with a 2 (contextual) × 2 (categorical) within-subjects design. Trials featuring stimuli from the same shape-sharing set (e.g., rolling pin with baguette/log/shovel) were excluded because they did not occur equally across conditions (i.e., they were absent from the within-within condition, see Experimental Design and Paradigm in methods for details). Figure 4A shows the amplitude of the difference between contralateral and ipsilateral sides for each condition. The ANOVA (Figure 3D) showed a significant main effect of category (F [1,23] = 4.96, *p* =.036) and a significant interaction between contextual and categorical similarity (F [1,23] = 6.16, *p* =.021). Post-hoc t-test analyses with Holm correction showed that the N2pc was significantly reduced when the target and distractor shared the context or the category, compared to when they shared neither (*p*_Holm_ < .05, two-tailed), but there was no further reduction when the target and distractor shared both.

## Discussion

We investigated how contextual associations guide attention and influence search performance. Our EEG results show that contextual associations affect attentional allocation jointly with categorical similarity during the selection stage (N2pc, 235-285 ms after onset). Our behavioral results show that distractors that share contextual associations with the target impair search performance both independently and in concert with categorical and perceptual similarity.

Our findings confirm the hypothesis that contextual associations guide attention during visual search, evidenced by attenuated N2pc effects when distractors share the same scene context as the target. Many visual search theories propose that information of target-relevant features is accumulated in parallel and then determines how attention is allocated to potential targets (Treisman & Sato, 1990; Wolfe, 1994, 2007, 2021). These theories are largely based on evidence from experiments using conjunctions of perceptual features (e.g., color and orientation). The present results extend these models by demonstrating that attentional guidance concurrently arises from multiple features, encompassing perceptual features and semantic features like contextual associations and category membership. In previous work, Moores et al. (2003) suggested that such semantic features guide attention during visual search. However, in their study, semantic features were defined broadly and stimulus pairs encompassed multiple types of relationships: For example, pairs with high co-occurrence probabilities could lack a common contextual associations (e.g., tables and chairs often go together, but across many possible contexts) and pairs with semantic associations could have low co-occurrence probabilities (e.g., a cow and a glass of milk are semantically related but not typically seen together). Moreover, categorical and perceptual similarities were not systematically controlled for, making it difficult to disentangle different types of semantic features and perceptual features. In the current study, we objectively measured perceptual similarity between targets and distractors, orthogonally manipulated contextual and categorical relationships, and applied linear regression to analytically control for their contribution. This allows us to characterize how attention is differentially guided by contextual and categorical similarity.

Our N2pc results revealed that contextual associations interacted with categorical effects to modulate attentional allocation to the target. Specifically, the N2pc amplitude decreased when distractors shared the same context or category as the target, with no additional reduction when both overlapped, indicating that contextual and categorical information jointly modulate attentional selection at this selection stage. The N2pc component likely indexes the final stage of attentional selection based on a priority map, where multiple guidance sources are combined in a weighted manner (Wolfe, 1994, 2021; Fecteau & Munoz, 2006; Eimer, 2014). The absence of an additive effect when both semantic relationships are present might be because contextual and categorical information are integrated within the priority map by selecting the feature that more efficiently discriminates between the target and the distractor (Yu et al., 2023). Although contextual and categorical relationships were manipulated in a binary manner in the present study, their relative strength is unlikely to be equal for each stimulus pair. For example, a cooking pot and a hand saw perhaps are more easily distinguished by context (kitchen vs. garden) than by category (tool vs. non-tool), indicating that the dominant semantic cue may depend on the specific object pair. Further, we found no interaction between perceptual similarity and either contextual or categorical effects in our EEG results. This contrasts with previous findings of a hierarchical interaction between perceptual and categorical effects during the N2pc window (Yeh & Peelen, 2022). A possible explanation is that feature-based guidance is task-dependent and adaptively relies on features that effectively distinguish the target from distractors (Wolfe, 2021; Yu et al., 2023; Yeh et al., 2024). In the previous study, perceptual similarity was relatively constrained: aside from distractors sharing an identical overall shape (e.g., round), the target was consistently paired with a distractor of a fixed, distinct shape (e.g., triangular). Under such conditions, perceptual features likely served as a very effective and often dominant cue for guiding attention. By contrast, in the present study, target-distractor perceptual similarity varied rather continuously across trials, given the stimuli were paired across six shape sets. Additionally, categorical similarity in this study was restricted to the superordinate level (tool vs. non-tool). Given that object categorization more commonly occurs at the basic level (e.g., a grater rather than a tool; Greene & Fei-Fei, 2014; Potter & Hagemann, 2014), future research should systematically investigate how different levels of category information interact with contextual associations and perceptual features to guide attention.

In addition to the EEG findings, we also found robust effects of contextual associations on search behavior. Specifically, targets and distractors sharing contextual associations impaired search accuracy both independently and in interaction with categorical similarity. The effect of contextual associations was stronger when distractors belonged to the same category as the target. By contrast, their influence on reaction time depended on perceptual similarity: the higher the perceptual similarity between the target and the distractor, the larger the contextual effect. Consistent with previous reports (Yeh & Peelen, 2022), we also observed main effects of perceptual and categorical similarity, as well as their interaction, on both accuracy and reaction time; greater perceptual similarity enhanced the categorical effect. The behavioral interactions of semantic and perceptual similarities are not found in the N2pc responses. Yet, behavioral performance reflects not only attentional guidance and selection but also later processes such as object identification and decision-making (Eimer, 2014; Wolfe, 2021). The observed interaction may therefore emerge at a later stage of processing, when the target template becomes more precise and incorporates detailed feature information (Wolfe, 2021; Yu et al., 2023).

Our study examined contextual associations based on object co-occurrence within a specific scene context (Bar, 2004; Oliva & Torralba, 2007). Here, we assumed that such co-occurrence statistics are relatively comparable across objects. Yet, this is clearly a simplification. First, objects may asymmetrically guide attention, where one object efficiently guides attention to the other, but not as much vice versa. Indeed, visual search in real-world scenes often relies on bigger “anchor objects” (e.g., sink, table) that guide attention when searching for associated smaller objects (e.g., toothbrush, mug) (Mack & Eckstein, 2011; Boettcher et al., 2018; Võ et al., 2019; Yu et al., 2023; Lerebourg et al., 2024; Souza-Wiggins & Geng, 2026). Second, effective attentional guidance may also depend on the typical relative positioning of objects, where attention is guided in directions that promise to be useful in the real world. For example, stoves guide attention upwards much more than downwards, reflecting the positioning of stove-associated objects above them (Kaiser et al., 2019; Torralba et al., 2006; Wolfe et al., 2019). How attentional guidance by contextual associations interacts with the asymmetries and spatial biases in real-world search remains an interesting open question for future research.

## Conclusion

The present study demonstrates that contextual associations work jointly with categorical information to guide attention, subsequently combining with both categorical and perceptual information to shape overall performance. These findings highlight that contextual associations support efficient attentional guidance in complex visual environments (e.g., searching for a cooking pot in a flea market). The context in which an object typically appears thus rapidly and robustly shapes visual cognition.

## Competing interest

No conflicts of interest, financial or otherwise, are declared by the authors.

## Acknowledgements

The authors thank Sirine Nouira for help with data collection in Experiment 2 and the same-different task.

## Funding

LCY is supported by the MSCA programme (101149060). DK is supported by the DFG (SFB/TRR135, project number 222641018; KA4683/5-1, project number 518483074, KA4683/6-1, project number 536053998) and an ERC Starting Grant (PEP, ERC-2022-STG 101076057). This work is further supported by the DFG under Germany’s Excellence Strategy (EXC 3066/1 “The Adaptive Mind”, project number 533717223). Views and opinions expressed are those of the authors only and do not necessarily reflect those of the funders. Neither the funders nor the granting authority can be held responsible for them.

## Data and materials availability

All study materials supporting this research are publicly available: https://osf.io/dwksa

## Supplementary

### Behavioral results

In Experiment 1, the error rate results showed significantly positive beta estimates for contextual similarity (b = .043, t[15] = 3.55, FDR-corrected *p* =.002), categorical similarity (b = .088, t[15] = 5.48, FDR-corrected *p* <.001), and perceptual similarity (b = 0.151, t[15] = 8.25, FDR-corrected *p* <.001), indicating that the higher the similarity, the worse the search accuracy. We also observed two significant interactions. The interaction between contextual and categorical similarity (b = .040, t[15] = 2.29, FDR-corrected *p* =.022) indicates that the contextual similarity effect was stronger when categorical similarity increased, and the interaction between categorical and perceptual similarity (b = .081, t[15] = 4.01, FDR-corrected *p* =.001) indicates that the categorical similarity effect was stronger when perceptual similarity increased. There was no interaction between contextual and perceptual similarity (b = .022, t[15] = 1.62, FDR-corrected *p* =.063). The response time results showed significantly positive beta estimates for categorical similarity (b = .060, t[15] = 3.70, FDR-corrected *p* =.003) and perceptual similarity (b = 0.124, t[15] = 5.33, FDR-corrected *p* <.001), indicating that the higher the similarity, the slower the response. We found no contextual effect (b = − 0.009, t[15] = −0.67, FDR-corrected *p* =.823) and no interactions (Contextual* Categorical: b = −0.012, t[15] = −0.95, FDR-corrected *p* =.823; Contextual* Perceptual: b = 0.009, t[15] = 0.66, p =.387; Categorical* Perceptual: b = 0.029, t[15] = 1.68, FDR-corrected *p* =.114).

In Experiment 2, the error rate results showed significantly positive beta estimates for contextual similarity (b = .027, t[23] = 2.70, FDR-corrected *p* =.009), categorical similarity (b = .074, t[23] = 7.61, p <.001), and perceptual similarity (b = 0.132, t[23] = 6.81, FDR-corrected *p* <.001), indicating that the higher the similarity, the slower the response. We also observed an interaction between categorical and perceptual similarity (b = .055, t[23] = 3.04, FDR-corrected *p* =.001), indicating that the categorical similarity effect was stronger when perceptual similarity increased. There were no significant interactions between contextual and categorical similarity (b = .010, t[23] = 0.78, FDR-corrected *p* =.265), and between contextual and perceptual similarity (b = .002, t[23] = 0.14, p =.446). The response time results showed significantly positive beta estimates for categorical similarity (b = .047, t[23] = 4.31, p <.001) and perceptual similarity (b = 0.125, t[23] = 7.93, FDR-corrected *p* <.001), indicating that the higher the similarity, the slower the response. No significant contextual effect was found (b = −.006, t[23] = −0.69, FDR-corrected *p* =.851). We also observed significant interactions between contextual and perceptual similarity (b = .050, t[23] = 3.22, FDR-corrected *p* =.003) and between categorical and perceptual similarity (b = .034, t[23] = 4.15, FDR-corrected *p* <.001), indicating that the contextual and categorical similarity effects were stronger when perceptual similarity increased. No significant interaction between contextual and categorical similarity was found (b = −.011, t[23] = −1.07, FDR-corrected *p* =.851).

## Notes

https://osf.io/dwksa

### Competing Interest Statement

The authors have declared no competing interest.

### Summary of Updates

1.The title has been changed to: “Contextual associations of real-world objects guide attention in visual search.” 2.The N1pc results have been removed in accordance with previous peer-review feedback.

## References

Alexander, R. G., & Zelinsky, G. J. (2012). Effects of part-based similarity on visual search: The Frankenbear experiment. Vision research, 54, 20–30.

Bahle, B., Winsler, K., Kiat, J. E., & Luck, S. J. (2025). Combined conceptual and perceptual control of visual attention in search for real-world objects. *Attention, Perception*, & Psychophysics, 1–19.

Bar, M. (2004). Visual objects in context. Nature Reviews Neuroscience, 5(8), 617–629.

Barras, C., & Kerzel, D. (2017). Target-nontarget similarity decreases search efficiency and increases stimulus-driven control in visual search. *Attention, Perception*, & Psychophysics, 79(7), 2037–2043.

Belke, E., Humphreys, G. W., Watson, D. G., Meyer, A. S., & Telling, A. L. (2008). Top-down effects of semantic knowledge in visual search are modulated by cognitive but not perceptual load. Perception & psychophysics, 70(8), 1444–1458.

Biederman, I. (1972). Perceiving real-world scenes. Science, 177(4043), 77–80.

Boettcher, S. E., Draschkow, D., Dienhart, E., & Võ, M. L. H. (2018). Anchoring visual search in scenes: Assessing the role of anchor objects on eye movements during visual search. Journal of vision, 18(13), 11–11.

Chun, M. M. (2000). Contextual cueing of visual attention. Trends in cognitive sciences, 4(5), 170–178.

Cornelissen, T. H., & Võ, M. L. H. (2017). Stuck on semantics: Processing of irrelevant object-scene inconsistencies modulates ongoing gaze behavior. *Attention, Perception*, & Psychophysics, 79(1), 154–168.

Desimone, R., & Duncan, J. (1995). Neural mechanisms of selective visual attention. Annual review of neuroscience, 18(1), 193–222.

Duncan, J., & Humphreys, G. W. (1989). Visual search and stimulus similarity. Psychological review, 96(3), 433.

Eimer, M. (1996). The N2pc component as an indicator of attentional selectivity. Electroencephalography and clinical neurophysiology, 99(3), 225–234.

Eimer, M. (2014). The neural basis of attentional control in visual search. Trends in cognitive sciences, 18(10), 526–535.

Fecteau, J. H., & Munoz, D. P. (2006). Salience, relevance, and firing: a priority map for target selection. Trends in cognitive sciences, 10(8), 382–390.

Greene, M. R., & Fei-Fei, L. (2014). Visual categorization is automatic and obligatory: Evidence from Stroop-like paradigm. Journal of vision, 14(1), 14–14.

Henare, D. T., Tünnermann, J., Wagner, I., Schütz, A. C., & Schubö, A. (2024). Complex trade-offs in a dual-target visual search task are indexed by lateralised ERP components. Scientific Reports, 14(1), 22839.

Jacob, G., & Arun, S. P. (2020). How the forest interacts with the trees: Multiscale shape integration explains global and local processing. Journal of Vision, 20(10), 20–20.

Jiang, Y., & Chun, M. M. (2001). Selective attention modulates implicit learning. The Quarterly Journal of Experimental Psychology: Section A, 54(4), 1105–1124.

Lleras, A., Wang, Z., Ng, G. J. P., Ballew, K., Xu, J., & Buetti, S. (2020). A target contrast signal theory of parallel processing in goal-directed search. *Attention, Perception*, & Psychophysics, 82(2), 394–425.

Luck, S. J., & Hillyard, S. A. (1994). Electrophysiological correlates of feature analysis during visual search. Psychophysiology, 31(3), 291–308.

Lerebourg, M., de Lange, F. P., & Peelen, M. V. (2024). Attentional guidance through object associations in visual cortex. Science Advances, 10(41), eado6226.

Mack, S. C., & Eckstein, M. P. (2011). Object co-occurrence serves as a contextual cue to guide and facilitate visual search in a natural viewing environment. Journal of vision, 11(9), 9–9.

Mirman, D., Landrigan, J. F., & Britt, A. E. (2017). Taxonomic and thematic semantic systems. Psychological bulletin, 143(5), 499.

Moores, E., Laiti, L., & Chelazzi, L. (2003). Associative knowledge controls deployment of visual selective attention. Nature neuroscience, 6(2), 182–189.

Nah, J. C., Malcolm, G. L., & Shomstein, S. (2021). Task-irrelevant semantic properties of objects impinge on sensory representations within the early visual cortex. Cerebral Cortex Communications, 2(3), tgab049.

Nah, J. C., & Geng, J. J. (2022). Thematic object pairs produce stronger and faster grouping than taxonomic pairs. Journal of Experimental Psychology: Human Perception and Performance, 48(12), 1325.

Nako, R., Wu, R., Smith, T. J., & Eimer, M. (2014). Item and category-based attentional control during search for real-world objects: Can you find the pants among the pans?. Journal of Experimental Psychology: Human Perception and Performance, 40(4), 1283.

Nako, R., Wu, R., & Eimer, M. (2014). Rapid guidance of visual search by object categories. Journal of Experimental Psychology: Human Perception and Performance, 40(1), 50.

Oliva, A., & Torralba, A. (2007). The role of context in object recognition. Trends in cognitive sciences, 11(12), 520–527.

Oostenveld, R., Fries, P., Maris, E., & Schoffelen, J. M. (2011). FieldTrip: open source software for advanced analysis of MEG, EEG, and invasive electrophysiological data. Computational intelligence and neuroscience, 2011(1), 156869.

31. Palmer, E. M., Van Wert, M. J., Horowitz, T. S., & Wolfe, J. M. (2019). Measuring the time course of selection during visual search. *Attention, Perception*, & Psychophysics, 81(1), 47–60.

Potter, M. C., & Hagmann, C. E. (2015). Banana or fruit? Detection and recognition across categorical levels in RSVP. Psychonomic Bulletin & Review, 22(2), 578–585.

Proulx, M. J., & Egeth, H. E. (2006). Target-nontarget similarity modulates stimulus-driven control in visual search. Psychonomic Bulletin & Review, 13(3), 524–529.

Seidl-Rathkopf, K. N., Turk-Browne, N. B., & Kastner, S. (2015). Automatic guidance of attention during real-world visual search. *Attention, Perception*, & Psychophysics, 77(6), 1881–1895.

Sisk, C. A., Remington, R. W., & Jiang, Y. V. (2019). Mechanisms of contextual cueing: A tutorial review. *Attention, Perception*, & Psychophysics, 81(8), 2571–2589.

Souza-Wiggins, M., & Geng, J. J. (2026). Anchor objects guide spatial attention during visual search. Attention, Perception, & Psychophysics, 88(1), 19.

Telling, A. L., Kumar, S., Meyer, A. S., & Humphreys, G. W. (2010). Electrophysiological evidence of semantic interference in visual search. Journal of Cognitive Neuroscience, 22(10), 2212–2225.

Torralba, A., Oliva, A., Castelhano, M. S., & Henderson, J. M. (2006). Contextual guidance of eye movements and attention in real-world scenes: the role of global features in object search. Psychological review, 113(4), 766.

Treisman, A., & Sato, S. (1990). Conjunction search revisited. Journal of experimental psychology: human perception and performance, 16(3), 459.

Võ, M. L. H., Boettcher, S. E., & Draschkow, D. (2019). Reading scenes: How scene grammar guides attention and aids perception in real-world environments. Current opinion in psychology, 29, 205–210.

Wolfe, J. M. (1994). Guided search 2.0 a revised model of visual search. Psychonomic bulletin & review, 1(2), 202–238.

Wolfe, J. M., & Gray, W. (2007). Guided search 4.0. Integrated models of cognitive systems, 99–119.

Wolfe, J. M. (2021). Guided Search 6.0: An updated model of visual search. Psychonomic bulletin & review, 28(4), 1060–1092.

Wu, C. C., Wick, F. A., & Pomplun, M. (2014). Guidance of visual attention by semantic information in real-world scenes. Frontiers in psychology, 5, 54.

Wyble, B., Folk, C., & Potter, M. C. (2013). Contingent attentional capture by conceptually relevant images. Journal of experimental psychology: human perception and performance, 39(3), 861.

Yeh, L. C., Yeh, Y. Y., & Kuo, B. C. (2019). Spatially specific attention mechanisms are sensitive to competition during visual search. Journal of Cognitive Neuroscience, 31(8), 1248–1259.

Yeh, L. C., & Peelen, M. V. (2022). The time course of categorical and perceptual similarity effects in visual search. Journal of Experimental Psychology: Human Perception and Performance, 48(10), 1069.

Yeh, L. C., Thorat, S., & Peelen, M. V. (2024). Predicting cued and oddball visual search performance from fMRI, MEG, and DNN neural representational similarity. Journal of Neuroscience, 44(12).

Yeh, L. C., Peelen, M. V., & Kaiser, D. (2026). Spatiotemporal representations of contextual associations for real-world objects. The Journal of Neuroscience, 46(19), e1967252026.

Yu, X., Zhou, Z., Becker, S. I., Boettcher, S. E., & Geng, J. J. (2023). Good-enough attentional guidance. Trends in Cognitive Sciences, 27(4), 391–403.

